# Accuracy of robotic coil positioning during transcranial magnetic stimulation

**DOI:** 10.1101/641456

**Authors:** Stefan M. Goetz, I. Cassie Kozyrkov, Bruce Luber, Sarah H. Lisanby, David L. K. Murphy, Warren M. Grill, Angel V. Peterchev

## Abstract

**Objective:** Robotic positioning systems for transcranial magnetic stimulation (TMS) promise improved accuracy and stability of coil placement, but there is limited data on their performance. This text investigates the usability, accuracy, and limitations of robotic coil placement with a commercial system, ANT Neuro, in a TMS study.

**Approach:** 21 subjects underwent a total of 79 TMS sessions corresponding to 160 hours under robotic coil control. Coil position and orientation were monitored concurrently through an additional neuronavigation system.

**Main Results:** Robot setup took on average 14.5 min. The robot achieved low position and orientation error with median 1.34 mm and 3.48°. The error increased over time at a rate of 0.4%/minute for both position and orientation.

**Significance:** After the elimination of several limitations, robotic TMS systems promise to substantially improve the accuracy and stability of manual coil position and orientation. Lack of pressure feedback and of manual adjustment of all coil degrees of freedom were limitations of this robotic system.

## Introduction

Coil placement has to be maintained during long stimulation sessions in the majority of research and clinical applications of transcranial magnetic stimulation (TMS). Conventional holders to stabilize the heavy stimulation coil are cumbersome, show elasticity of their arms as well as joints, render adjustments with millimeter accuracy difficult, and do not compensate for subject movement [1, 2]. Alternatively, manually holding the coil throughout long sessions requires continuous attention, causes operator fatigue, depends on operator skill, and is costly [3].

Several navigated robotic positioning systems for TMS have been introduced to address these shortcomings [2, 4-9]. These systems include a robotic arm that holds the coil in a specified position and orientation relative to the subject’s head. With the aid of sensors, such as stereocameras, the coil placement is maintained automatically even if the subject moves [7]. Given the growing need for precision in coil placement to match the development of fMRI-guided targeting, robotic systems show great promise. However, there is a lack of information about the performance of such systems in routine TMS studies.

This paper reports on the accuracy and user experience with an ANT Neuro robotic positioning system during real-world conditions of a TMS study.

## Methods

We used a robotic system with adaptive positioning (ANT Neuro, Enschede, Netherlands) including a six-axis industrial robot (Omron Adept Viper s650, San Ramon, CA) and a tracking camera (NDI Polaris, Waterloo, Canada). We monitored and recorded the position and orientation of the coil at the time of each stimulus with six degrees of freedom (three spatial coordinates and three angles) using a second stereotactic system (Brainsight, Rogue Research, Montreal, Canada). The second stereotactic system received a trigger from the stimulator to synchronize recording of position and orientation with the pulse (Figure 1).

**Figure 1.**
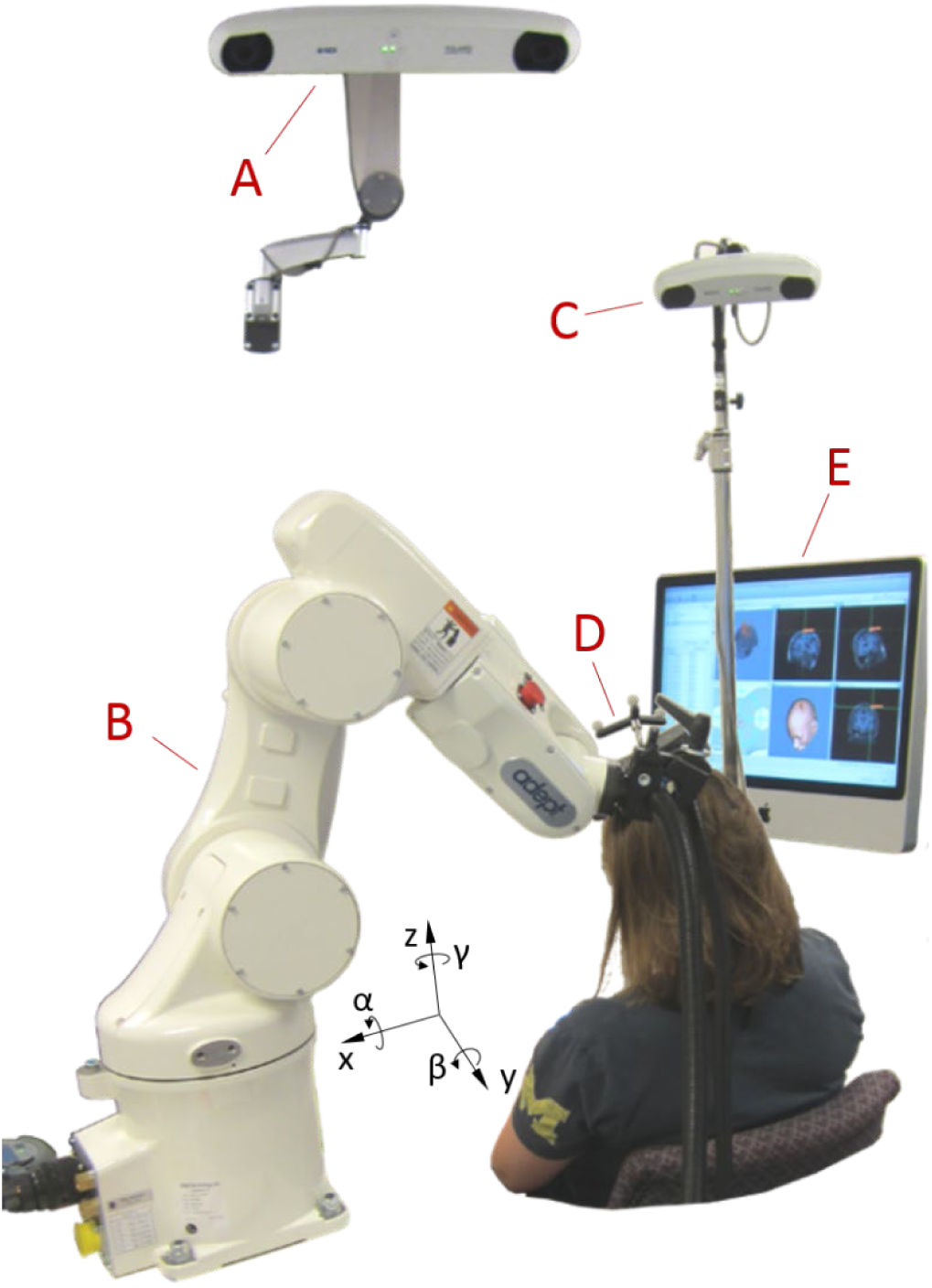
Picture of the robotic TMS setup, showing a stereocamera for the robot control system (A), an industrial robot (B), a stereocamera for monitoring the coil placement (C), a TMS coil with tracker (D), and a neuronavigation system for monitoring the coil placement (E). All position recordings are relative to the head (Talairach coordinates); therefore the x direction refers to the axis pointing from right to left, y from anterior to posterior, and z from inferior to superior, while α denotes the angle of rotation around the x axis (pitch), β around the y axis (yaw), and γ around the z axis (roll).

The coil positioning data were obtained during experiments for a previously published repetitive TMS study [11]. TMS was performed on 21 subjects (age 18 – 48 years, median 21, 14 females, 7 males, all right handed) in 1–7 sessions (3.76 sessions per subject on average). Sessions lasted 48–194 min (mean 124 min), and contained both single-pulse TMS with jittered inter-stimulus intervals between 8 s and 12 s and 1 Hz repetitive TMS over the left primary motor cortex. In total, 106,904 coil position samples synchronized to stimulation pulses were recorded.

Subjects were seated in a comfortable chair and told to move their head as little as possible and only very slowly so that the robot could compensate for movement. The two stereotactic systems each used its own camera but the same reflective fiducial markers on the subject’s head, also called optical trackers in the following (Figure 1). For both stereotactic systems, each subject’s head was registered using characteristic anatomic landmarks (nasion, right and left intertragic notches) identified by the same operator throughout the study. The same operator also performed the coil and robot setup in every session. For the orientation control and collision prevention of the robot, the surface of the head was sampled with at least 800 points covering the entire surface formed by the frontal, occipital, parietal, and temporal bones, including at least 300 points in the target area. For tracking of head position, we used glasses with reflective trackers and double-sided tape on the nasal bridge to stabilize the position of the glasses throughout the session. For safety, we avoided any head rest or fixation as the robot could press the subject’s head against it. The center of the figure-of-eight coil (Magventure Cool-B65, Farum, Denmark) was covered with a layer of compressible gauze, generating light pressure and friction for comfort and stabilization of the coil–head contact.

Statistical analysis was performed in JMP (SAS Institute, Cary, NC, USA). Prior to further effect screening, all data underwent analysis for statistical distribution; exclusively nonnegative data was also subjected to Box-Cox distribution analysis based on Akaike’s Information Criterion (AIC). With appropriate data transformation to incorporate the specific statistical distribution, we analyzed the absolute position and orientation and the deviation from the position and orientation of the location found to generate largest MEPs (“hotspot”) with mixedeffects models.

From the time course of the errors of position and orientation, we extracted the time the robot required after major deflections to reposition the coil again and correct 80% of the deviation in position or orientation. Major deflections were defined as exceeding the error of the previous sample by at least 20% and 2 mm or 5° for the position and orientation, respectively.

## Results

The net setup time of the robot for a subject at the beginning of each session was (14.48 ± 4.22) min (mean ± standard deviation), defined as the time from the last neuronavigated handheld hotspotting pulse to the first robot-controlled pulse.

The use of the robot led to low targeting errors throughout the entire study with a median of 1.34 mm and 3.48° for position and orientation, respectively. The various effects contributing to the error are summarized in Figure 2 and Table 1.

**Table 1.** Results of mixed effects model of the robot’s absolute position and deviation from the target. Significant effects (p < 0.05) are highlighted in bold.

| Variable | Factor | Statistics | Value |
| --- | --- | --- | --- |
| Absolute position & orientation* | Sex | **F(1,106904) = 528, p < 0.001** | for females position shifted by<br>x: -2.17 mm<br>y: -5.15 mm<br>z: -2.83 mm |
|  | Age | **F(1,106904) = 3290, p < 0.001** | shift per year of age<br>x: +0.149 mm<br>y: -0.382 mm<br>z: -0.387 mm |
| | Session time | F(1,106904) = 0.0031, p = 0.956 | -27.1 $\mu\text{m}/\text{min}$ in x,<br>-141 $\mu\text{m}/\text{min}$ in y,<br>+6.88 $\mu\text{m}/\text{min}$ in z,<br>-0.0233°/min in $\alpha$,<br>-0.0400°/min in $\beta$,<br>+0.0331°/min in $\gamma$ |
|  | Session number | F(6, 106904) = 1.91, p = 0.076 |  |
| | Sex × Session time | F(1,106904) = 0.159, p = 0.690 | |
| | Age × Session time | $F(1,106904) = 2.96, p = 0.0852$ | |
|  | **Session time × Session number** | **$F(6,106904) = 5.32, p < 0.001$** |  |
| Position error<sup>†</sup> | **Sex** | **$F(1,106904) = 96.8, p < 0.001$** | 6.82% higher for females |
|  | **Age** | **$F(1,106904) = 2170, p < 0.001$** | −1.76% per year of age |
|  | **Session time** | **$F(1,106904) = 577, p < 0.001$** | 0.404% per minute |
|  | **Session number** | **$F(6,106904) = 4811, p < 0.001$** |  |
|  | **Sex × Session time** | **$F(1,106904) = 467, p < 0.001$** | 0.172% per minute for females, 0.643% per minute for males |
|  | **Age × Session time** | **$F(1,106904) = 102, p < 0.001$** | −0.119% rate decrease per year of age |
|  | **Session number × Session time** | **$F(6,106904) = 258, p < 0.001$** |  |
| Orientation error<sup>††</sup> | **Sex** | **$F(1,106904) = 5634, p < 0.001$** | 65.2% larger for males |
|  | **Age** | **$F(1,106904) = 8814, p < 0.001$** | −3.52% per year |
|  | **Session time** | **$F(1,106904) = 526, p < 0.001$** | 0.387% per minute |
|  | **Session number** | **$F(6,106904) = 2523, p < 0.001$** |  |
|  | **Age × Session time** | **$F(1,106904) = 66.4, p < 0.001$** | −0.0964% rate decrease per year of age |
|  | **Session number × Session time** | **$F(6,106904) = 769, p < 0.001$** |  |
| | **Sex × Session time** | $F(1,106904) = 1.32, p = 0.2958$ | 0.376% per minute for males, 0.401% per minute for females |
| Duration of position correction after major deflection<sup>‡</sup> | **Sex** | $F(1,79) = 1.09, p = 0.301$ | +17.8% for males |
| | **Age** | $F(1,79) = 1.75, p = 0.190$ | +1.2% per year of age |
| | **Session number** | $F(6,79) = 1.53, p = 0.192$ | |
| Duration of orientation correction after major deflection<sup>‡‡</sup> | **Sex** | $F(1,79) = 1.39, p = 0.238$ | +18.9% for males |
|  | **Age** | **$F(1,79) = 5.10, p = 0.027$** | 1.9% per year of age |
| | **Session number** | $F(6,79) = 1.81, p = 0.102$ | |

\* normal distribution of position ( $AIC(normal) = 9.21 \cdot 10^5$ , $AIC(Weibull) = 9.60 \cdot 10^5$ , $AIC(lognormal) > 1.5 \cdot 10^6$ ) and orientation ( $AIC(normal) = 3.26 \cdot 10^5$ , $AIC(Weibull) = 4.80 \cdot 10^5$ , $AIC(lognormal) > 9.0 \cdot 10^5$ )
<sup>†</sup> lognormal distribution ( $AIC(normal) = 1.02 \cdot 10^6$ , $AIC(Weibull) = 4.23 \cdot 10^5$ , $AIC(lognormal) = 3.39 \cdot 10^5$ , Box-Cox Coefficient = −0.1)
<sup>††</sup> lognormal distribution ( $AIC(normal) = 5.44 \cdot 10^5$ , $AIC(Weibull) = 4.79 \cdot 10^5$ , $AIC(lognormal) = 4.70 \cdot 10^5$ , Box-Cox Coefficient = 0.2)
<sup>‡</sup> lognormal distribution ( $AIC(normal) = 612$ , $AIC(Weibull) = 528$ , $AIC(lognormal) = 494$ , Box-Cox Coefficient = −0.25)
<sup>‡‡</sup> lognormal distribution ( $AIC(normal) = 487$ , $AIC(Weibull) = 436$ , $AIC(lognormal) = 417$ , Box-Cox Coefficient = −0.051)

**Figure 2.**
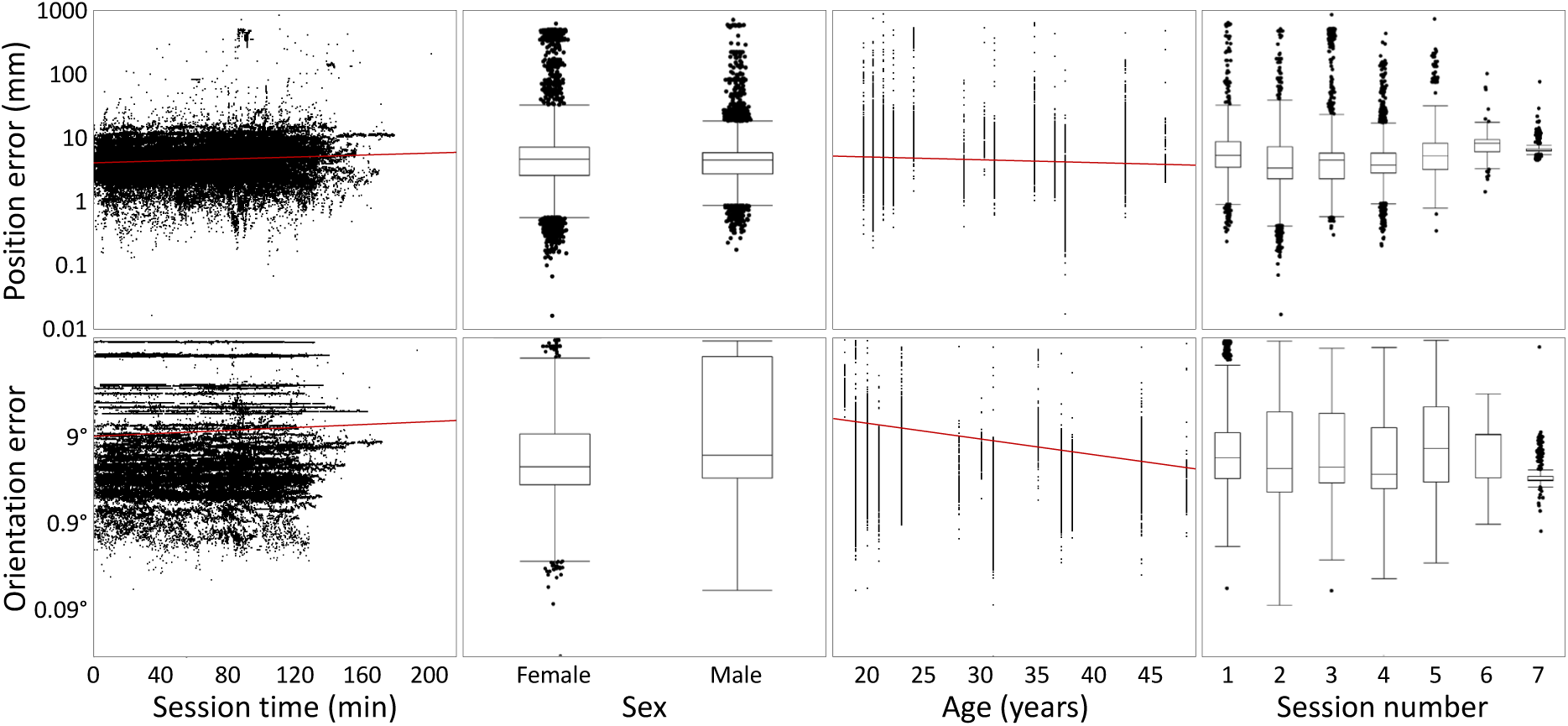
Dependence of the position error (top) and the orientation error (bottom) on key effects. Red lines indicate regression trends on the logarithmic scale.

The absolute position depended on Sex and Age, which may have influenced height as well as the seating position. There was also an interaction between Session number and Session time. Importantly, the effect of Session time was insignificant, indicating that the coil placement did not drift systematically in any direction.

The error of the coil position with respect to the target identified at the beginning of each session, was on average (3.9 ± 28.0) mm (mean ± standard deviation) (see Figure 2). The orientation error was on average (4.04 ± 3.13) ° (mean ± standard deviation). The correlation coefficient between position and orientation error was linearly 0.594 (p < 0.001) and logarithmically 0.179 (p < 0.001).

The deviation of the position from the target was dependent on Sex, Age, Session time, Session number, and the interactions of Sex and Session time, of Age and Session time, and of Session number and Session time. The overall position error decreased with age, while older subjects showed a slower increase of the error.

Likewise, the orientation error depended on Sex, Age, Session time, Session number, and the interactions of Age and Session time and of Session number and Session time (see Table 1). The error grew over time with 0.387% per minute. Further, both the overall orientation error as well as the speed of increase of the orientation error decreased with age.

Large brief targeting errors up to 860 mm occurred after subjects rapidly moved their head away from the coil. After such major deflections, the time for the robot to return to baseline was (10.7 ± 11.4) s (mean ± standard deviation). The duration for orientation errors to return to baseline was (8.14 ± 7.05) s (mean ± standard deviation). The durations for both the return of the position and orientation exhibited approximately lognormal distributions. For both position and orientation errors, age and male sex tended to increase the duration for correction, and the effect of age was significant for orientation errors.

## Discussion

In this robotic rTMS study, we detected coil position errors consistent with the maximum values of 2 mm reported in previous benchtop measurements of similar systems [5, 8]. The identified repositioning error of 3.64 mm between sessions of the same subject is likely less an indicator of robot error than of different head registration errors between the sessions. The robot is comparable or may outperform the accuracy of manual placement with stereotactic support, which was reported to range from 2 mm to more than 5 mm, although our study did not intend to compare robotic and manual coil placement [12, 13]. The practicality of robotic placement appears superior for long sessions, in which the additional setup time of about 15 min is acceptable. A conclusive quantitative comparison with manual placement concerning positioning accuracy, on the other hand, will require a one-to-one comparison with a repeated measures setup, which appears justified as soon as the revealed technical limitations are solved.

The overall error and to a lesser extent the rate of increase of the position and angular errors were smaller for older individuals. These trends may be related to the anecdotally observed increased compliance of older subjects with the guideline to keep the head as still as possible. Frequent head movements can lead to constant errors at the time of pulses as the robot repositioning lags behind and to growing errors because of movement-mediated shift of the head trackers.

While the absolute position was not dependent on the session time, the deviation from the target was. Accordingly, the error grew over time, but the robot did not show a preferential absolute direction. Thus, all three directions contributed approximately similarly to deviations.

The positioning errors can be attributed to several factors. The repeatability of positioning of industrial robots is typically excellent, even though coil position and orientation are estimated exclusively from the states of the six joints along the kinematic chain [14]. Therefore, the largest portion of the error may result from control. The controller estimates the local head curvature at the target from an idealized head scaled to the sampled outline of the subject. This procedure typically does not consider individual differences of the head shape and may misrepresent the surface curvature and the normal direction [15].

With a misestimated local curvature, the coil does not contact the head with its focus point, which is approximately in the center of the bottom face of a figure-of-eight coil. Thus, the coil is not perfectly tangential to the head at the focus point. This issue is often underestimated in manual coil positioning as well, where it requires experienced operators to appreciate and manage the impact of imperfect tangential orientation on the head surface. For example, on a sphere with a radius of 85 mm, which represents a typical head curvature at the motor cortex, rolling the coil by only 5° shifts the point on the scalp where the coil touches the head by 7.4 mm and lifts the focal point of the coil by 0.65 mm, further increasing the distance to the target [16-18].

The ANT robot control software allowed corrections of the three translational degrees of freedom as well as the orientation of the coil relative to the central gyrus. However, it did not allow manual adjustment of the two angular degrees of freedom that control the position of the coil surface relative to the assumed local head surface normal. Whereas one of the two degrees of freedom can be and was corrected by rolling the coil handle in the clamp, the other one did not allow a simple mechanical adjustment. It is expected that corrections of less than 5° could control a large portion of the constant part of the observed positioning error.

The current study also revealed that the coil position constantly drifts relative to the target without any preferential direction so that targeting accuracy decreases over time. On average, however, the rate of drift appears moderate on the order of 0.4% per minute (i.e., a doubling time of about 175 min) for both position and orientation so that an error of 1 mm and 5° increases to 1.27 mm and 6.3° over an hour. This growing offset may arise from a number of contributions. Dominant may be minor movements and position drifts of the optical trackers on the coil and the subject, which are a key limitation of all frameless stereotaxy systems in TMS [13, 19]. These drifts accumulate over time and are amplified by movements of the subject. Tracker movements can deteriorate the tight coil–head contact, without which subjects tend to constantly reposition their head and the robot lags behind causing a deviation. In contrast to widely-used expandable headbands, which can easily move and allow the lever that accommodates the trackers to swing, we used trackers on glasses worn by the subject. The position of the glasses is comparably well defined as they rest on the rigid nasal bone and their orientation is stabilized by the temples which provide a long lever to the ears. However, despite our use of double-sided tape to fix the eyeglass bridge to the skin, movement of the skin underneath the bridge can still contribute to the error.

Whereas this study used a commercial system based on an industrial robot with a relatively moderate cost, other purpose-designed robots have been presented or are commercially available for TMS [5, 7, 9, 20]. The alternative robot mechanics are unlikely to improve positioning accuracy, as the accuracy is limited by other factors, and indeed bench measurements showed deviations that are comparable with industrial robots [9]. Nevertheless, kinematics that are adjusted to the typical movement range of the human head can improve repositioning speed and range, which is limited with small industrial robots. During larger rotation or movement of the head that would require the robot to leave its range, the robot risks losing the target entirely, which we observed several times during training sessions.

A major issue was identified prior to the study. The coil surface abutting the head is smooth and rigid. Without some pressure against the coil, subjects tended to perform constant minor position adjustments of the head, which the robot compensated with time-lagged millimeter-scale oscillatory movements. Accordingly, the robotic system would require a highly accurate adjustment of the coil– head contact in 0.1 mm steps to find an equilibrium between the following two cases as observed in our lab: In one scenario, the robotically actuated coil slowly pushed the subject’s head into uncomfortable positions and potentially out of range throughout the session, which can happen with speeds as slow as a few millimeters per minute. In the opposite scenario, there was a loose (no pressure) mechanical contact between the coil and the head. In this case the subject may lean into the coil, to which the robot responds by moving the coil away, again resulting in a slow drift.

This contact issue is a direct consequence of the robot control approach, which is exclusively based on position. For this study, we introduced passive pressure control as soon as the focal point of the coil touched the head by locally adding compressible gauze between the coil and the head. For small coil–scalp distances, the gauze was compressed to a thin layer and the gauze elasticity converted the coil–scalp distance into contact pressure. Thus, if the target location is set for a position where the coil compresses the gauze tightly but not entirely, an increase of the pressure by the subject moves the coil away, whereas a release attracts the coils, keeping the relative position to the target and the pressure practically constant. In addition, the increased friction between the scalp and the coil reduces the small continuous oscillatory movements. The pressure control and the additional friction turn out to be better controllable for the application of TMS coil positioning.

Robot systems that provide force control based on pressure sensors promise better control over the coil–head interface [9]. Multi-cell pressure sensors can further ensure that the coil touches the target with its focus point and is tangential to the head surface at the target [10].

## Acknowledgements

Research reported in this publication was supported by the Duke-Coulter Translational Partnership, Brain & Behavior Research Foundation under NARSAD Young Investigator Award 22796, and National Institutes of Health under award numbers RF1MH114268 and U01AG050618, and in part by the Intramural Research Program of the National Institute of Mental Health (ZIAMH00295). The content is solely the responsibility of the authors and does not necessarily represent the official views of the National Institutes of Health. We thank Dr. L. Gregory Appelbaum and Dr. Lysianne Beynel for their valuable comments on the manuscript.

S. M. Goetz, S. H. Lisanby, D. L. K. Murphy, and A. V. Peterchev are inventors on patents and patent applications on TMS technology. S. M. Goetz has received research funding from Magstim Inc. A. V. Peterchev has received research and travel support as well as patent royalties from Rogue Research; research and travel support, consulting fees, as well as equipment donation from Tal Medical/Neurex; patent application and research support from Magstim; as well as equipment loans from MagVenture, all related to TMS technology. I. C. Kozyrkov, B. Luber, and W. M. Grill report no relevant disclosures.

